# Amyloid fibril structure of islet amyloid polypeptide by cryo-electron microscopy reveals similarities with amyloid beta

**DOI:** 10.1101/2020.02.11.944546

**Authors:** Christine Röder, Tatsiana Kupreichyk, Lothar Gremer, Luisa U. Schäfer, Karunakar R. Pothula, Raimond B. G. Ravelli, Dieter Willbold, Wolfgang Hoyer, Gunnar F. Schröder

## Abstract

A critical role of the hormone islet amyloid polypeptide (IAPP) is vividly discussed for Type 2 Diabetes (T2D), where amyloid deposits in pancreatic islets consisting of fibrillar IAPP have been associated with beta cell loss. Here, we applied cryo-electron microscopy to elucidate the structure of IAPP fibrils prepared at physiological pH and reconstructed densities of three dominant polymorphs. An atomic model of the main polymorph comprising residues 13 – 37 in a density map of 4.2 Å resolution reveals two S-shaped, intertwined protofilaments. The segment 21-NNFGAIL-27, which is essential for IAPP amyloidogenicity, forms the protofilament interface together with tyrosine 37 and the amidated C-terminus. The main IAPP fibril polymorph resembles polymorphs of the Alzheimer disease (AD)-associated amyloid-β peptide (Aβ), which is striking in light of the epidemiological link between T2D and AD and reports on IAPP-Aβ cross-seeding *in vivo*. The results structurally link the early-onset T2D IAPP genetic polymorphism S20G with the early-onset AD Arctic mutation E22G of Aβ, rationalize previous data on IAPP fibrils, help to elucidate mechanisms of amyloid formation and toxicity, and support the design of fibril growth inhibitors as well as imaging probes for early detection of IAPP fibrils.

## Introduction

Pancreatic islet amyloid deposits are a hallmark of type 2 diabetes (T2D). Islet amyloid, first reported almost 120 years ago as islet hyaline^1^ is found in > 90 % of individuals with T2D^2,3^. The main constituents of islet amyloid are fibrillar aggregates of the 37-residue polypeptide hormone islet amyloid peptide (IAPP), also called amylin. IAPP is detected in many organs including the brain but is mainly localized in the beta cells of the pancreatic islets, where it is co-synthesized and co-secreted with insulin^3,4^. IAPP is involved in glucose homeostasis and metabolism, with putative functions as regulator of insulin and glucagon secretion, satiety, and gastric emptying^3,5^. Formation of toxic IAPP amyloid aggregates has been associated withdysfunction and death of beta cells, placing T2D in the group of protein misfolding disorders^2,3,5–8^. However, the nature of the toxic IAPP species and the mechanisms of beta cell death are not well determined^9^. Potential toxic effects of IAPP amyloid comprise induction of apoptosis^10^, chronic inflammation^11^, defects in autophagy^12,13^, endoplasmic reticulum stress^14,15^, and membrane disruption^16^. Apart from its association with T2D, IAPP amyloid might also play a role in type 1 diabetes^10,17^.

IAPP interacts with amyloidogenic proteins that trigger other protein misfolding disorders^18–20^. Of particular interest is its relation to the amyloid-β peptide (Aβ) peptide, the main component of the senile plaques found in Alzheimer’s disease (AD) brains. IAPP and Aβ are not only both infamous for their strong aggregation propensity and the insolubility of their aggregates^3^, but they also display primary sequence similarity^21^. IAPP and Aβ colocalize in islet amyloid in T2D patients and in Aβ deposits in brains of AD patients^19^. Mutual cross-seeding of IAPP and Aβ aggregation observed in transgenic mice further supports a role of the IAPP-Aβ interaction in pathogenesis^19,20^.

Structural information on IAPP amyloid is essential as a basis for improving our understanding of the mechanism of amyloid formation, for defining toxic IAPP species, and for elucidating IAPP-Aβ cross-seeding^5,7^. Furthermore, high-resolution IAPP fibril structures can inform the design of fibril growth inhibitors and support the development of soluble, nontoxic IAPP analogs for coformulation with insulin or leptin for therapy of type 1 diabetes or obesity, respectively^5^. Current structural models of IAPP fibrils at physiological pH, based for example on solid-state NMR of full-length IAPP and X-ray crystallography of IAPP fragments, consistently place the majority of the 37 amino acid residues into the fibril core while the N-terminus is located at the periphery^22–27^. On the other hand, the available models also exhibit substantial differences, which could either be a consequence of the limited, distinct restraints obtained by the different techniques applied or a reflection of IAPP fibril polymorphism^5,28^. Here, we have applied cryo-electron microscopy (cryo-EM) to determine the structure of IAPP amyloid fibrils grown at physiologically relevant pH. We provide a structural analysis of three dominant polymorphs, including an atomic model of the main polymorph comprising residues 13 – 37 in a density map of 4.2 Å resolution.

## Results

For this work, amyloid fibrils were prepared from synthetic human IAPP (hIAPP) including the amidated C-terminus. Islet amyloid in T2D is typically extracellular, but IAPP aggregation is supposedly initiated intracellularly, possibly in the secretory granules at a pH of 5.0 – 6.0^3,29^. IAPP fibrils were therefore prepared at pH 6.0. Long, well-ordered fibrils were obtained as shown by AFM imaging (Supplementary Figures 1 and 2). We could differentiate at least five different polymorphs in the AFM images and in subsequently performed cryo-EM experiments. Of these five polymorphs, three have been present in sufficient amounts for further analysis (Figure 1, Supplementary Figures 1 and 2). The main polymorph, polymorph 1 (PM1) makes up ∼90 % of all fibrils, while polymorph 2 (PM2) and polymorph 3 (PM3) might represent up to ∼10 % and ∼1 %, respectively, of the total number of fibrils in the dataset. The structure and molecular details of these three polymorphs (Figure 1) are further described in the following sections.

**Figure 1.**
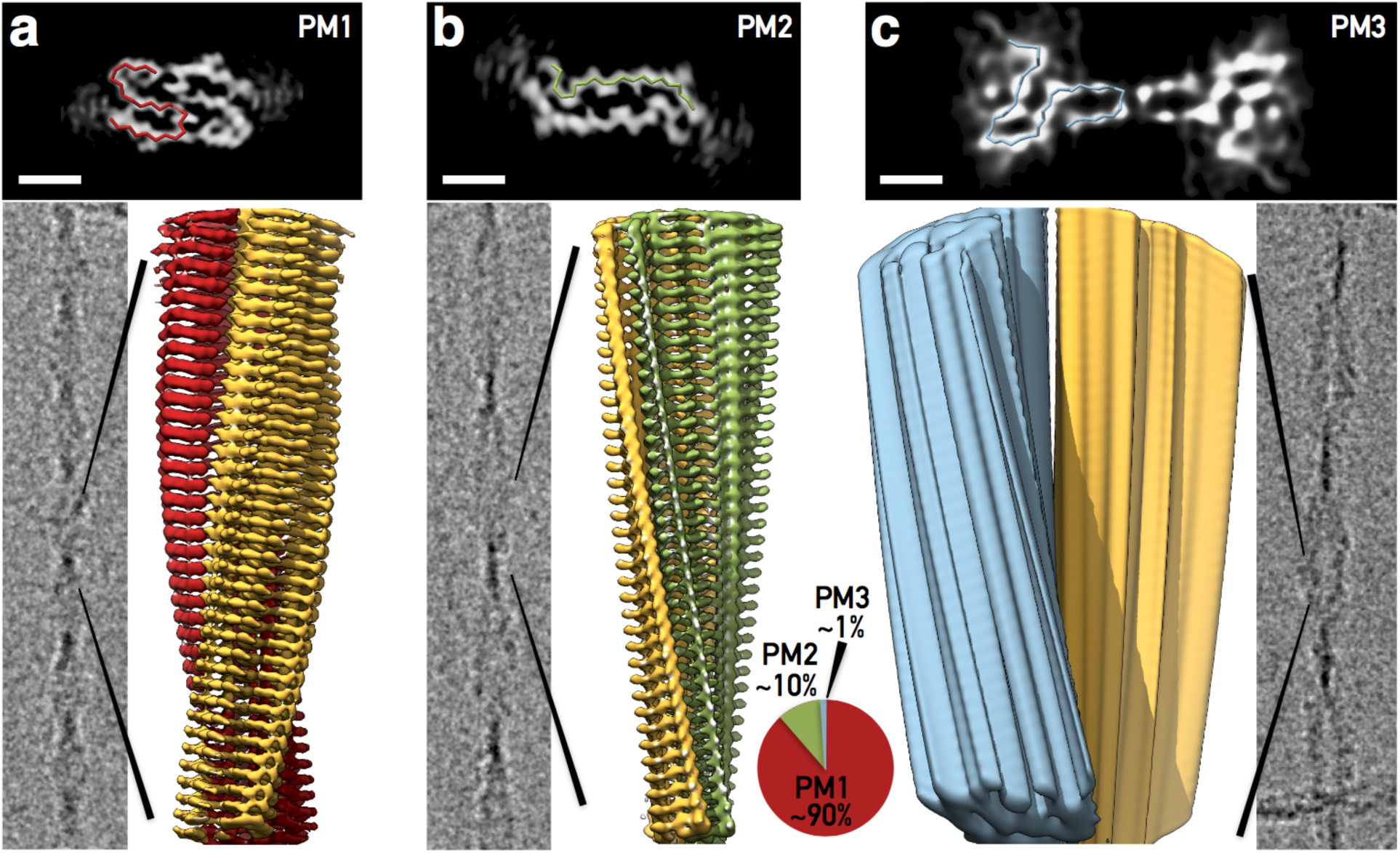
Comparison of reconstructed IAPP polymorphs. PM1 (**a**), PM2 (**b**), and PM3 (**c**). For each polymorph three panels are shown: a slice of a 3D reconstruction superimposed with the respective Cα-chain for one monomer (top, black box, scale bars are 2 nm), a micrograph displaying the respective polymorph (grey) and a three-dimensional density (PM1, red/yellow (**a**); PM2, green/yellow (**b**); PM3, blue/yellow (**c**)). The pie chart visualizes the fraction of each polymorph in the data set.

### Architecture of polymorph 1

The most dominant polymorph (PM1) has a right-handed helical symmetry with a pitch of 48 nm and a width of 2.5 – 4.5 nm (Figures 1 a, 2 a). 3D reconstruction of 1161 individual fibril images using a pseudo 2_1_-symmetry led to a resolution of 4.2 Å, which was sufficient to unambiguously build an atomic model (Figure 2 b) with the helical parameters being 2.35 Å (helical rise) and 178.23° (helical twist). The fibril consists of two stacks of IAPP monomers winding around each other. Each monomer exhibits an overall S-fold that clearly displays residues Ala^13^ – Tyr^37^ packed into three β-sheets (Figure 3 a). Up to residue 12, the N-terminal part including the disulfide bond between Cys^2^ and Cys^7^ is largely disordered and therefore does not reveal clear density (Figure 1 a). The side view of PM1 shows the typical cross-β pattern of amyloid fibril structures with a spacing of 4.7 Å between the layers. The cross-β layers are well resolved in the density as shown in Figure 2 d. On the secondary structure level, we observed three β-sheets: residues 14 – 20, 26 – 32, and 35 – 37. Figure 3 shows the comparison of our model with former secondary structure predictions based on sequence analysis^22^, NMR^23,26,27^, EPR^25^, and X-Ray experiments^24^.

**Figure 2.**
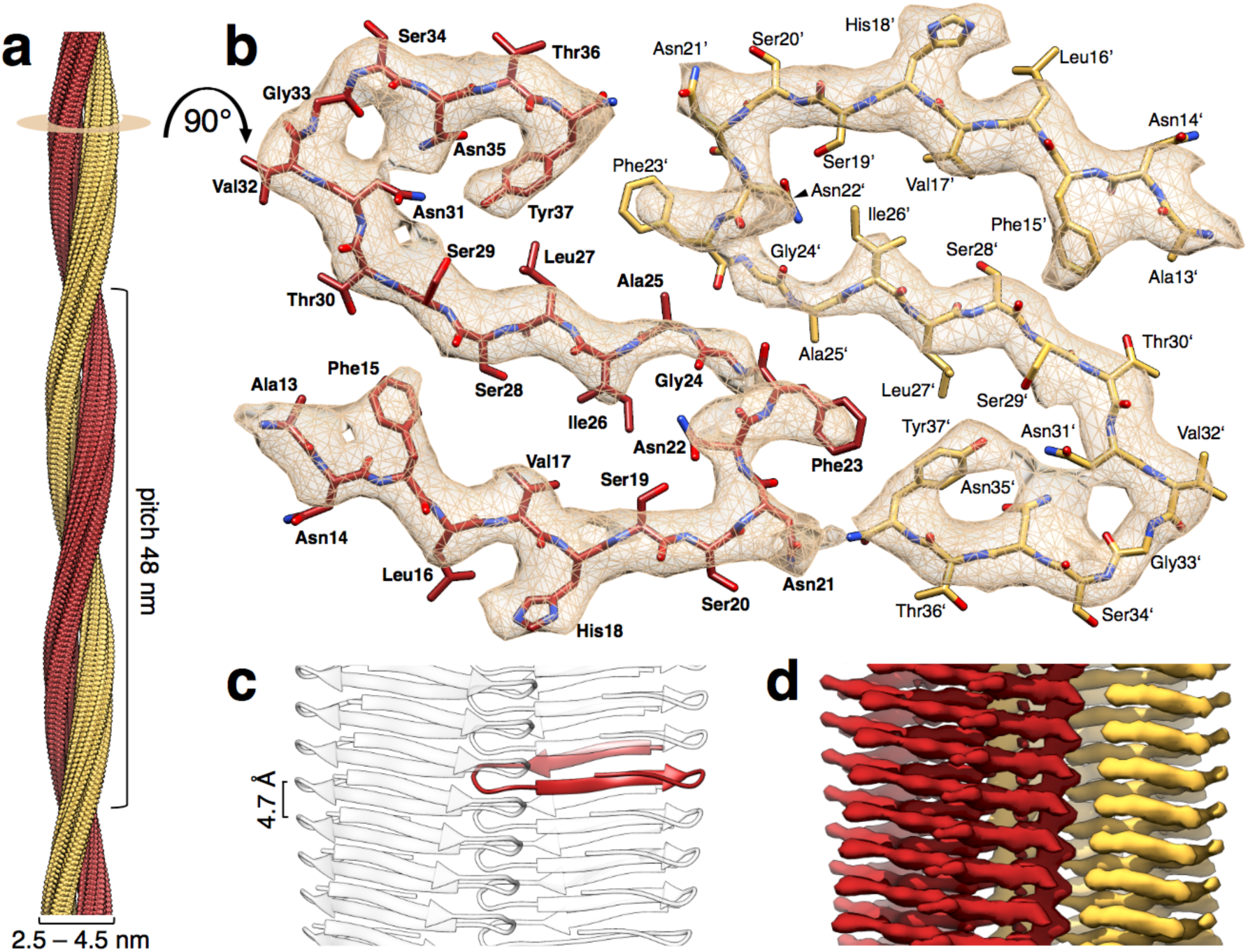
Structural details of the main polymorph PM1. **a** PM1 exhibits a helical pitch of 48 nm and a minimum and maximum width of 2.5 and 4.5 nm, respectively. The fibril consists of two protofilaments (red and yellow). **b** The cross-section displays two symmetry-related monomers with an atomic model of residues 13 – 37 built into the 4.2 Å density (contour level of 1.5 σ). **c** Side view of the fibril model. One monomer is highlighted (red) to show its integration in the fibrillar structure. Cross-β layers are separated by 4.7 Å. **d** Side view of the reconstructed density corresponding to (**c**).

**Figure 3.**
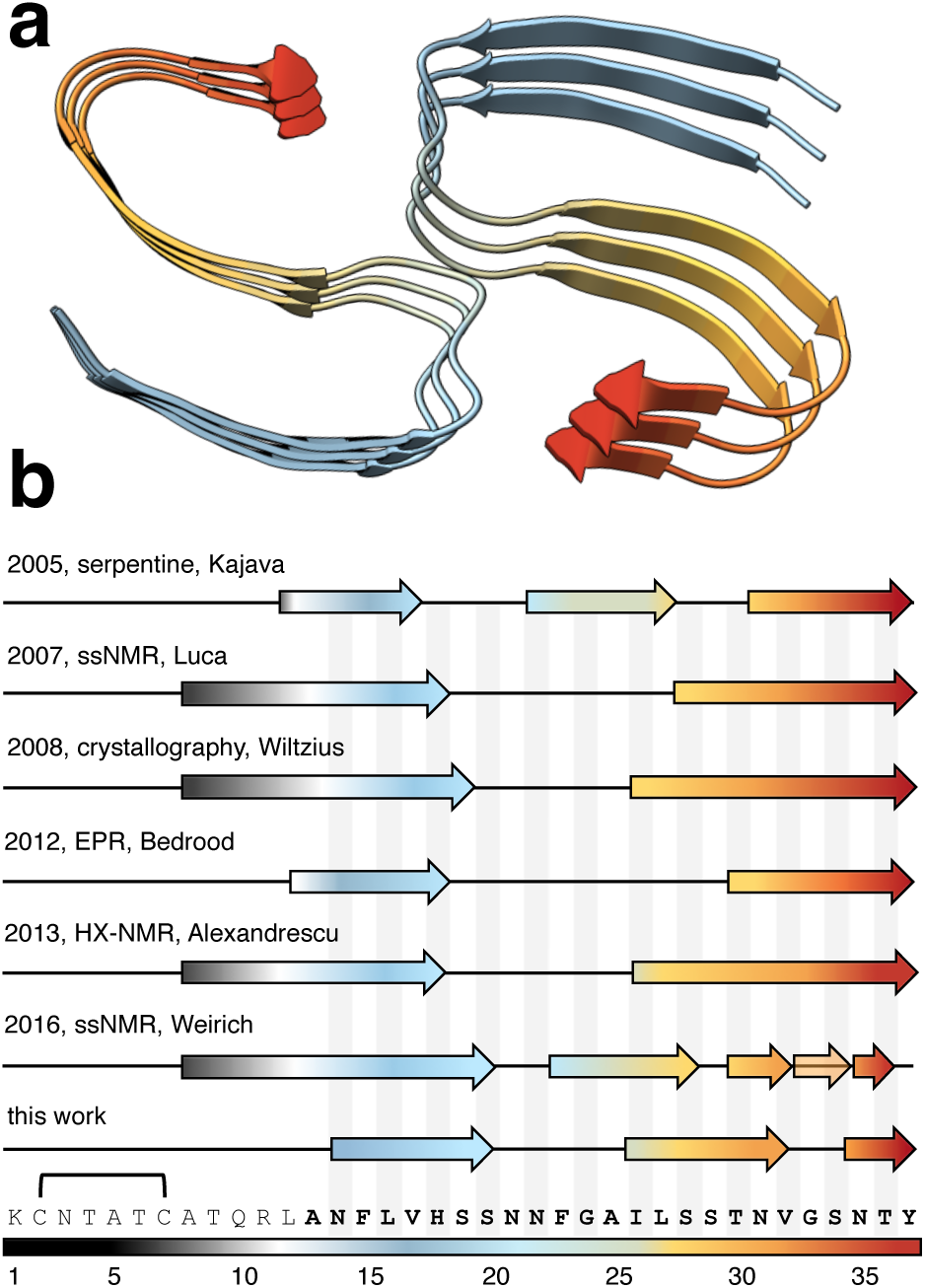
Secondary structure comparison of PM1 with previous IAPP fibril models. **a** Secondary structure of the IAPP model presented in this study. Tilted cross-section of three fibril layers with representation of the three β-sheets observed. **b** Comparison of our PM1 structure to former models from sequence-based prediction^22^, EPR^25^, ssNMR^27^, X-ray crystallography^24^, and hydrogen-exchange (HX) NMR studies^26^. Arrows indicate β-sheets (one potential β-sheet is shown as transparent). The disulfide bridge between residues Cys^2^ and Cys^7^ is indicated. Fibril formation was performed at pH 7.4^23,25–27^ or pH 6.5^24^, while our IAPP fibrils were formed at pH 6.0.

The cross section of the PM1 fibril displays two monomeric S-folds related by the approximate 2_1_-symmetry (Figure 2 a). The double-S shape is stabilized both by hydrophobic and polar interactions. The central part of the protofilament interface consists of a hydrophobic cluster comprising residues Phe^23^, Gly^24^, Ala^25^, Leu^27^ and Phe^23’^, Gly^24’^, Ala^25’^ Leu^27’^ (Figure 2b and Supplementary Figure 3). Additionally, the backbone of Phe^23^ and Ala^25^ forms hydrogen bonds at the center of the fibril (Figure 4 a) thereby connecting one subunit with two neighboring subunits above and below in the opposing protofilament (Figure 4 b). More precisely, there is a hydrogen bond between carbonyl-Phe^23^ of chain *i* and amide-Ala^25^ of chain *i*+1 and another hydrogen bond between amide-Ala^25^ of chain *i* and carbonyl-Phe^23^ of chain *i*-1. The backbone around Gly^24^ does not maintain the cross-β hydrogen bonding pattern along the fibril. The aforementioned interactions are formed by residues located in the sequence motif (N)NFGAIL that has been shown earlier to be important for fibrillization of IAPP^5,30–32^. This motif is located in the central part of the structure in the turn between the first two β-sheets (Figures 2 and 3). Within this turn, the kink around Phe^23^ and Asn^21^ is stabilized by hydrogen bonds between Asn^22^ and Ser^19^ as well as between Asn^22^ and Gly^24^ (Figures 2 and 4). Additionally, Ile^26^ might support this turn by hydrophobic interactions with Val^17^. In the second turn – between β-sheet 2 and 3 – Asn^31^ together with Ser^29^, Asn^35^ and Tyr^37^ creates a hydrophilic cluster at the C-terminus of IAPP with possible interactions between Asn^31^ and Ser^29^ as well as Asn^31^ and Asn^35^. Tyr^37^ might as well interact with both, Asn^35^ and Ser^29^ (Figures 2 and 4). Moreover, the amidated C-terminus itself forms a polar ladder (Figure 4 b). This ladder is further connected to Asn^21’^ of the opposite protofilament with slightly longer and therefore weaker hydrogen bonds (Figure 4). The overall cross-β arrangement is further stabilized by Asn^14^, Asn^21^, and Asn^31^, which form polar ladders alongside the fibril axis. Asn^22^ does not form a polar ladder, but instead its Nδ2 atom forms a hydrogen bond to the carbonyl-Gly^24^ within the same monomer (Figure 4 a).

**Figure 4.**
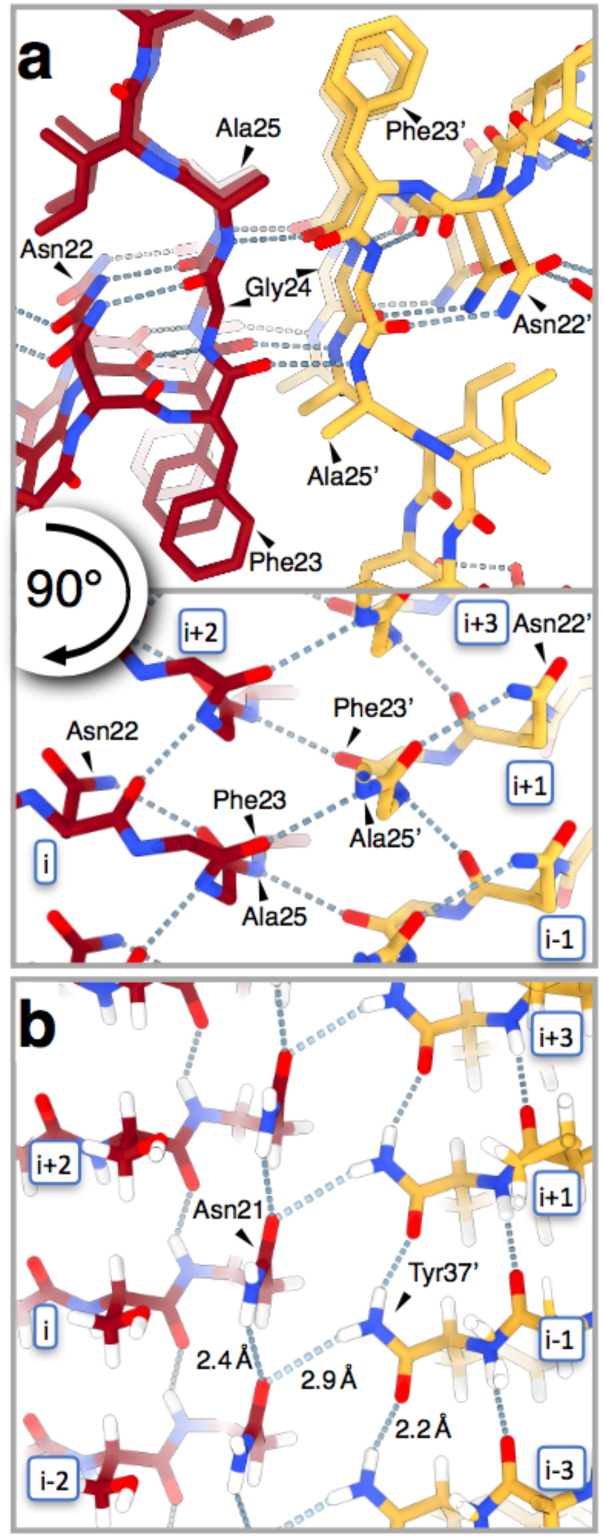
Details of IAPP polymorph PM1. **a** Top view of the fibril core showing the two NFGAIL motifs in the opposing protofilaments (red, yellow) as well as the hydrogen bonding network (dashed lines). The panel below shows a side view of the fibril core and illustrates the interlocking of the protofilaments by hydrogen bonds. **b** Side view of the fibril showing the hydrogen bonding interaction of Asn^21^ with the amidated C-terminus of the opposing protofilament (yellow) as well as polar ladders of Asn^21^ and Tyr^37’^ along the fibril axis.

All-atom MD simulations were performed to evaluate the overall stability of the model. In two independent 250 ns simulations, the model remained stable (Supplementary Figure 5) with an all-atom root mean square deviation (RMSD) of a single subunit from the deposited model of ∼2 Å and a root mean square fluctuation (RMSF) of residues 16 – 37 of 0.8 Å. The N-terminal part including Phe^15^ is already significantly more mobile (Supplementary Figure 5 a, b, e). Notably, we do observe ladder formation for Asn^22^ in the MD simulation, which is not supported by the density map.

For earlier structures of amyloid fibrils we have discussed the need for a minimal fibril unit, which is the smallest fibril structure fragment in which the capping subunits at both ends would have established the same full contact interface with other constituting monomers as the capping subunits of an extended fibril^33,34^. Here, the minimal fibril unit consists of only three monomers, which is the smallest possible unit. One subunit is in contact exclusively with its neighboring monomers above and below and its opposing monomers through protofilament interface contacts (Figure 2 c). Indeed, we could not observe any sort of interlocking of different cross-β layers that was postulated to have a stabilizing effect on other amyloid fibrils^33,34^.

### Structure modeling for polymorph 2

Like PM1, polymorph 2 (PM2) also consists of two protofilaments and exhibits a 2_1_-symmetry (Figure 1 a, b). With a maximum and minimum width of 52 Å and 17 Å, respectively, PM2 shows a more pronounced twist in the projection images (Figure 1 b) and is remarkably flatter than PM1 (Figure 1 a). The helical pitch is 94 nm and AFM experiments suggest a left-handed twist. In contrast to the S-shaped PM1, the density map indicates an extended conformation of two IAPP monomers in PM2. The protofilament interface consists of a continuous sequence region of at least 18 amino acids. The density map with a resolution of about 4.2 Å would in principle allow for model building of 21 amino acid residues, however, the sequence assignment is ambiguous. We have therefore modeled all 17 possible sequence assignments both in forward and backward backbone trace directions, leading to 17*2=34 different models. All 34 models were refined in DireX^35^ using for cross-validation the resolution range of 3.0 – 4.0 Å for calculation of the C_free_ value^36^. Results were ranked by C_free_ value (Figure 5, Supplementary Figure 6). According to this criterion, the most probable model for PM2, which also exhibits the highest C_work_ value, shares important features with the PM1 model. First, the NFGAIL motif forms the center of the fibril interface. Second, the N-terminus is rather flexible and thus not resolved in the density map (Figure 5). The first visible residue in the density of this model of PM2 is Phe^15^. In contrast to PM1, not only the N-terminus, but also the two C-terminal residues Thr^36^ and Tyr^37^ are not clearly resolved and are potentially mobile. In between the two protofilaments is a relatively large cavity lined by hydrophobic residues Phe^23^, Ala^25^, and Ile^26^. It is not clear whether this gap is water filled.

**Figure 5.**
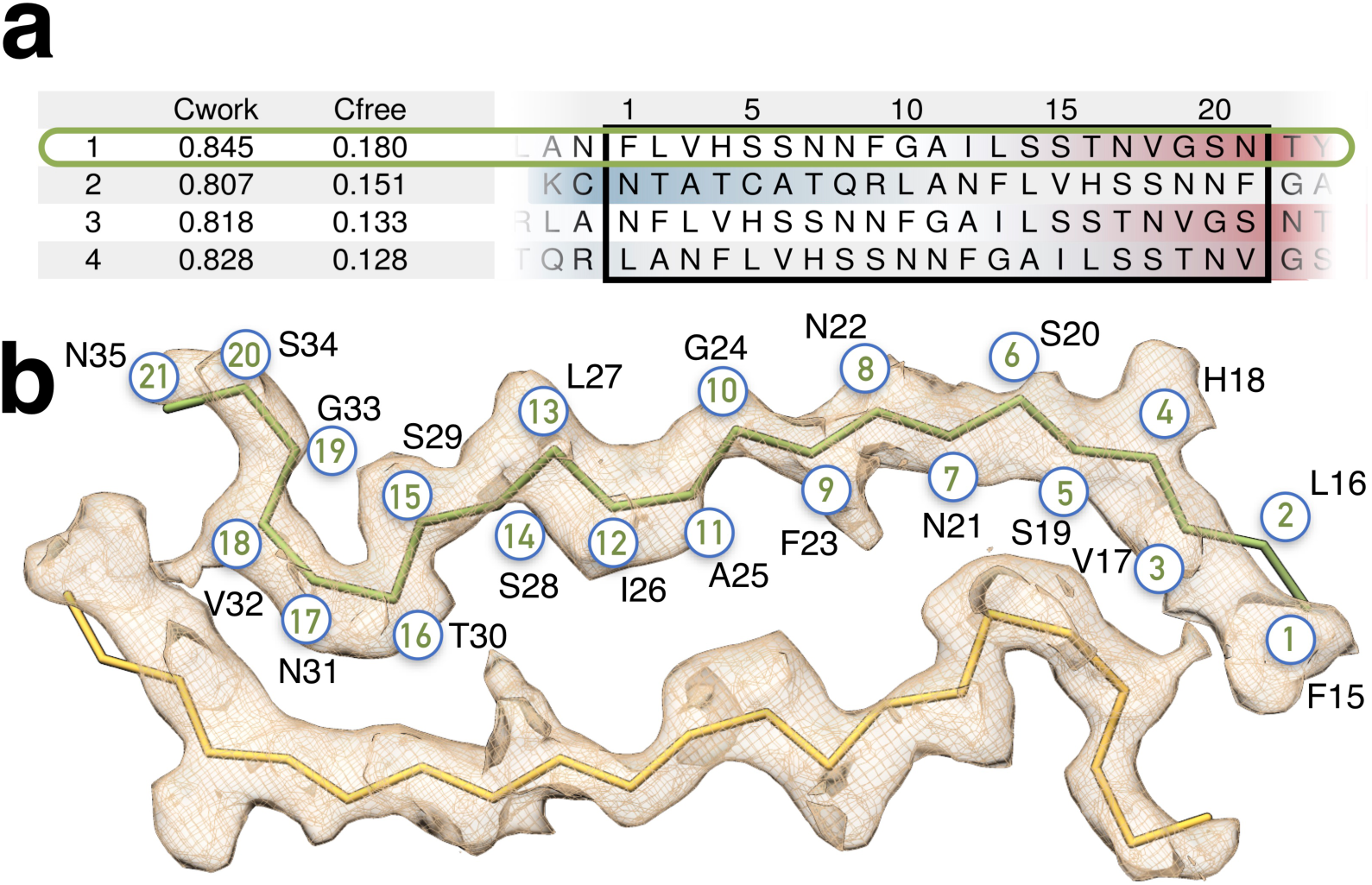
Results from structure fitting into the density of PM2. **a** The table contains the C_work_ and C_free_ values from DireX fitting of 21-residue-long sequence snippets (black box) of IAPP into a density layer of PM2 together with the respective amino acid sequence. Highlighted (green box) is the most favorable sequence fit (see also Supplementary Figure 6). **b** The cross-section of PM2 density with a Cα-trace. Encircled numbers represent the residue position according to (**a**) (black box). Residues Phe^15^ to Asn^35^ (black letters) visualize the most favorable model highlighted in (**a**).

### Polymorph 3 exhibits a narrow interface

Compared to the other polymorphs, polymorph 3 (PM3) was not well represented in the micrographs. The overall features of PM3, namely the broad width (110 Å) and pronounced twist (159 nm pitch), lead to a dumbbell shape (Figure 1 c). From the 4591 particles extracted, we could reconstruct a density with a resolution of 8.1 Å. Since the resolution is rather low, we were not able to build an atomic model, but only hypothesize a possible Cα backbone trace. Nonetheless, the density clearly indicates again two symmetric protofilaments (Figure 1) and displays that the 10 Å wide protofilament interface of PM3, presumably consisting of three residues, is very small compared to those of PM1 and PM2.

## Discussion

The IAPP fibril samples investigated here displayed fibril polymorphism, with the dominant polymorphs PM1, PM2, and PM3 accounting for ∼90%, ∼10%, and ∼1% of all fibrils, respectively. While all three polymorphs consist of two (pseudo-)symmetric helically intertwined protofilaments, they display substantial differences in the protein fold (Figure 1). PM1 consists of a compact S-shaped fold, PM2 features an extended IAPP conformation, and the PM3 cross-section shows two compact motifs connected by an extended bridge. Marked differences are also observed between the protofilament interfaces. In PM1, the interface consists of one of the turns and the C-terminal end of the S-fold. In PM2, the entire extended IAPP segment that constitutes the fibril core is involved in the protofilament interface. Finally, PM3 exhibits a very narrow interface of probably three residues only. Despite of these differences, certain IAPP sequence segments might contribute similarly to distinct fibril polymorphs: in both PM1 and the most probable PM2 model, residues 22-NFGAIL-27 form the central fibril core.

In an early report^28^ of IAPP fibril polymorphism the most common polymorph consisted of two protofilaments coiled around each other with a helical pitch of 50 nm, and another polymorph showed a helical pitch of 100 nm. These values are in good agreement with PM1 (48 nm) and PM2 (94 nm). Despite these similarities, when comparing the cryo-EM results with previous structural data it has to be kept in mind that differences may arise both from differences between the applied techniques and from differences between the polymorphs present in the samples. In agreement with previous studies^22–27^, we find that the IAPP N-terminus including the disulfide bond between Cys^2^ and Cys^7^ is not part of the fibril core, neither in PM1 (Figure 3 b) nor in the most probable model of PM2 (Figure 5). While well-defined density starts from residue 13 in the cryo-EM data, some studies reported the fibril core to begin around residue 8 (Figure 3 b). Hydrogen-exchange (HX) NMR data, however, indicated that residues 8-14 were less protected than those in the central fibril core^26^. In agreement with previous data, residues 13-37 are largely in β-sheet conformation in PM1, although there is variation with respect to the precise location of β-strands (Figure 3 b). A common feature of the PM1 cryo-EM structure and previous models is a turn in the segment 20-SNNFG-24^23–27^. A second turn is formed in PM1 in the segment 32-VGS-34. This turn was also supported by ssNMR and HX NMR studies^26,27^. The two turns establish an S-shaped fold of IAPP in PM1. As a consequence, the tyrosyl ring of the C-terminal Tyr^37^ packs against Phe^23’^ in the adjacent protofilament which is in line with distance restraints for IAPP fibrils obtained by fluorescence resonance energy transfer (FRET)^37^. In addition, FRET experiments proposed a maximum distance of 11 Å between Tyr^37^ and a second phenylalanine which is in agreement to the Tyr^37^-Phe^15^ distance in the PM1 model^37^. The C-terminal amide stabilizes the S-shaped fibril structure by forming a polar ladder and a hydrogen bond with Asn^21^ in the adjacent protofilament (Figure 4 b), in line with enhanced amyloid formation upon C-terminal amidation of IAPP^38,39^.

The sequence region 20 – 29 is particularly important for amyloidogenicity of IAPP^5,30,31^. This can be rationalized with the PM1 fibril structure: First, residues 22-NFGAILSS-29 constitute the solvent-excluded central core of PM1 fibrils (Figure 2 b). Second, residues 21-NNFGAIL-27 form, together with Tyr^37^ and the amidated C-terminus, the protofilament interface. In previous structural models of IAPP fibrils the region 20-29 was associated with formation of a partially ordered loop rather than a β-structure, which was surprising considering the sensitivity of IAPP amyloid formation to mutations in this region^7^. The PM1 fibril structure shows that residues 20-25 indeed form a turn, albeit one that is an integral part of the fibril core, featuring an extensive hydrogen bond network (Figure 4). Residues 26-29, on the other hand, are part of the central β-sheet of IAPP PM1 fibrils. Remarkably, the structure of the 21-NNFGAIL-27 segment in PM1 is highly similar down to the atomic detail to a crystal structure of the NNFGAIL peptide^24^ (Figure 6 a). This applies both to the fold of the individual polypeptide molecules and to the peptide/protofilament interface, which displays extensive mainchain-mainchain interactions between the 23-FGA-25 segments (Figure 6 a). The similarity of the NNFGAIL structure in the peptide crystal and in the PM1 fibril suggests that it is indeed the 21-NNFGAIL-27 segment that drives IAPP amyloid formation.

**Figure 6.**
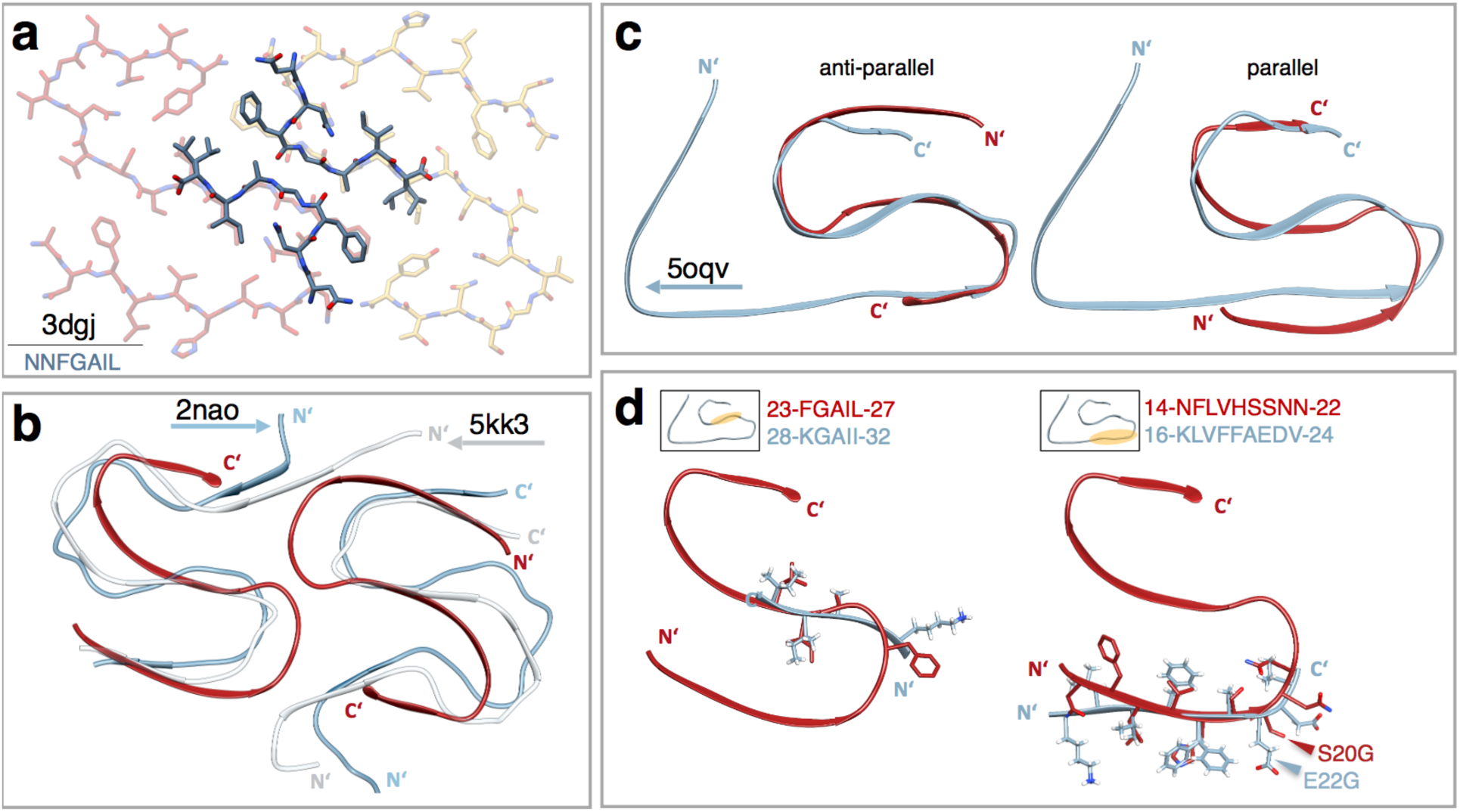
Structure comparison of PM1 with a fibrillar IAPP peptide (a) and amyloid-β fibril models. (**b-d**). **a** Overlay of our model with a crystal structure (dark) of the NNFGAIL peptide from Wiltzius *et al.*^24^. **b** Overlay of IAPP (red) with NMR structures by Waelti *et al.*^55^ (light blue, 2nao) and Colvin *et al.*^59^ (grey, 5kk3). **c** Anti-parallel (left) and parallel (right) overlay of one IAPP PM1 monomer (red) with the atomic model of Aβ(1-42) (light blue) from Gremer *et al.*^33^ **d** Detailed parallel overlay of sequence segments in IAPP (red) and Aβ(1-42) (light blue) from Gremer *et al.*^33^. Small boxes indicate where the respective sequence motif is located in the Aβ(1-42) model. The FGAIL motif of IAPP (red) shows high sequence identity to KGAII (Aβ, light blue). The NFLVHSSNN motif of IAPP (red) corresponds to the KLVFFAEDV motif of Aβ(1-42) with high structural similarity (right). Disease-related substitutions in IAPP (S20G) and Aβ(1-42) (E22G) are located at corresponding indicated positions.

IAPP contains an unusually large number of the polar residues asparagine, serine and threonine^5^. We find that these residues form polar streaks within the fibril core of PM1 and of the most probable model of PM2 (Figure 5, Supplementary Figure 6). The polar streaks are characterized by extensive networks of hydrogen bonds. The segregation of polar and apolar residues into distinct clusters within the fibril core likely contributes to the high stability of IAPP amyloid. In IAPP, the segregation is facilitated by the pre-organization of amino acid residues in polar and apolar clusters within the primary structure, in the fashion of a block copolymer with polar blocks 19-SSNN-22 and 28-SSTN-31 and the apolar block 23-FGAIL-27.

A serine-to-glycine substitution at position 20 (S20G), the only known IAPP genetic polymorphism in humans, is associated with early onset of T2D^40,41^. The S20G substitution enhances aggregation and toxicity of IAPP and leads to increased beta cell apoptosis^42–45^. Substitution of serine with glycine was suggested to promote turn formation at residue 20, favoring the amyloid fibril conformation^46,47^. In line with this notion, Ser^20^ is located at the edge of the turn comprising residues 20-25 in PM1. Interestingly, when comparing the S-fold of IAPP with the LS-fold of Aβ^33^ (Figure 6 c,d), the S20G substitution in IAPP and the Arctic mutation E22G of Aβ^48^, which causes early-onset AD, are located at corresponding positions (Figure 6 d). This suggests that these two replacements with glycine might have analog conformational consequences.

In contrast to the human protein, IAPP from several other species was found to be non-amyloidogenic^5^. The non-amyloidogenic rat and mouse IAPP contains six amino acids different from the human sequence^31^. Five of those are located in the sequence region 23–29 which is part of the central core of PM1 fibrils as discussed above. The differing amino acids include three prolines in rat and mouse IAPP at positions 25, 28 and 29. As proline disrupts secondary structures, these proline residues are incompatible with the PM1 structure, in line with the low amyloidogenicity of rat and mouse IAPP.

The insights gained from the rat and mouse IAPP sequences were exploited in the design of pramlintide, a non-amyloidogenic IAPP analog carrying proline substitutions at positions 25, 28 and 29^49^. Pramlintide is co-administered with insulin in type 1 diabetes to improve glucose level regulation. Similarly, combination of a non-amyloidogenic IAPP analog and leptin might be interesting for treatment of obesity^50^. However, these drugs would benefit from increased solubility^51^. The structural data on IAPP fibrils presented here may aid in the design of non-amyloidogenic, soluble IAPP analogs by suggesting potential sites for chemical modifications of IAPP that counteract fibril formation.

Colocalization of IAPP and Aβ has been observed in T2D and AD patients^19^. The epidemiological association of diabetes and dementia might be explained by cross-seeding of IAPP and Aβ aggregation^19,20,52,53^. Different sites on amyloid fibrils are relevant for cross-seeding: cross-elongation, i.e. the elongation of a fibril with a heterologous protein, occurs at the fibril end, while cross-nucleation, i.e. the fibril-catalyzed formation of a heterologous fibril nucleus, may occur both at the fibril end and along the fibril surface. Like IAPP, Aβ forms different fibril polymorphs according to ssNMR and cryo-EM studies ^33,54–58^. Comparing IAPP PM1 to multiple Aβ(1–42) polymorphs containing S-shaped folds^33,55,59^, we find that the backbones superimpose (Figure 6 b, c). The structural similarity of the backbones is highest when superimposing the models in an antiparallel arrangement (Figure 6 c). The similarity between IAPP and Aβ(1–42) fibril folds regarding topology and size might promote cross-seeding at the fibril end, which could further be supported by the sequence similarity of IAPP and Aβ^21^. The sequence similarity is highest around the Gly-Ala-Ile segment at position 24–26 of IAPP and position 29–31 of Aβ. In both IAPP and Aβ, this segment is located in the solvent-excluded center of the S-fold (Figure 6 d). A further segment that can be superimposed in parallel arrangement is the N-terminal strand of the S-fold in IAPP PM1 and in the LS-shaped Aβ(1–42) polymorph, 14-NFLVHSSNN-22 of IAPP and 16-KLVFFAEDV-24 of Aβ (Figure 6 d). As noted above, the S20G IAPP substitution and the E22G Arctic Aβ mutation are located at corresponding positions in this superposition.

This study presents the 4.2 Å structure of an IAPP fibril polymorph consisting of two S-shaped protofilaments, but also highlights the polymorphism of IAPP fibrils. The dominant S-shaped PM1 can rationalize many of the characteristics of IAPP fibrils described by various groups, suggesting that it is a common polymorph or that it at least represents general features of prevalent IAPP polymorphs. The study provides detailed insight into the link between the IAPP amino acid sequence and fibril structure. It furthermore reveals similarities between IAPP and Aβ fibril structures, which are particularly striking in consideration of the link between diabetes and AD. The structural information gained will serve as a basis to define the mechanisms of amyloid formation and toxicity of IAPP. Moreover, the PM1 fibril may be used as a target structure to design imaging probes for IAPP fibrils and inhibitors of IAPP fibril growth.

## Methods

### Sample preparation

Human IAPP (H-KCNTATCATQRLANFLVHSSNNFGAILSSTNVGSNTY-NH_2_, molecular mass 3903.4 Da) with amidated C-terminus and the disulfide bond between Cys^2^ and Cys^7^ was custom synthesized by Pepscan (Lelystad, Netherlands). Identity and purity (93.2%) were confirmed by RP-HPLC and mass spectroscopy. To ensure monomeric starting material, the peptide was dissolved at 2 mg·ml^-1^ in 1,1,1,3,3,3-hexafluoro-2-propanol at RT for 1 h and lyophilized. Afterwards, 1 mg peptide powder was dissolved in 0.5 ml aqueous 6 M guanidine hydrochloride solution and size exclusion chromatography was performed on a Superdex 75 Increase 10/300 column (GE Healthcare) equilibrated with 10 mM 2-(N-morpholino)ethanesulfonic acid (MES)/NaOH buffer pH 6.0 using an ÄKTA Purifier system (GE Healthcare). The monomeric peak fraction was collected, aliquoted, flash-frozen in liquid nitrogen and stored at -80°C for further use. IAPP fibrils were prepared from the stock solution by diluting to a final concentration of 100 µM peptide with 10 mM MES/NaOH buffer pH 6.0, 6 mM NaN_3_. Fibrillation occurred by incubation for 7 days at RT under quiescent conditions in 1.5 ml Protein LoBind tubes (Eppendorf).

### Atomic force microscopy (AFM)

IAPP fibrils in 10 mM MES/NaOH buffer pH 6.0, 6 mM NaN_3_ were diluted to a peptide concentration of 10 µM monomer equivalent. Afterwards, 5 µl of the fibril solution were applied onto freshly cleaved muscovite mica and incubated under a humid atmosphere for 10 minutes. After three washing steps with 100 µl ddH_2_O the samples were dried with a stream of N_2_ gas. Imaging was performed in intermittent contact mode (AC mode) in a JPK Nano Wizard 3 atomic force microscope (JPK, Berlin) using a silicon cantilever with silicon tip (OMCL-AC160TS-R3, Olympus) with a typical tip radius of 9 ± 2 nm, a force constant of 26 N/m and resonance frequency around 300 kHz. The images were processed using JPK DP Data Processing Software (version spm-5.0.84). For the presented height profiles a polynomial fit was subtracted from each scan line first independently and then using limited data range.

### Cryo-EM image acquisition

Cryo-preparation was performed on glow-discharged holey carbon films (Quantifoil R 1.2/1.3, 300 mesh). A total of 2.5 µl sample containing 100 µM IAPP in 10 mM MES, 6 mM NaN_3_, pH 6.0, was applied to the carbon grid and incubated for 1 min. Subsequently, the sample was blotted for 5 s (blotting force 5) before being cryo-plunged using a Vitrobot (FEI). With 110,000-fold nominal magnification 1330 micrographs have been recorded on a Tecnai Arctica electron microscope operating at 200 kV with a field emission gun using a Falcon III (FEI) direct electron detector in electron counting mode directed by EPU data collection software (version 1.5). Each movie was composed of 50 fractions. Each fraction contained 36 frames, i.e. a total of 1800 frames were recorded per micrograph. The sample was exposed for 46.33 s to an integrated flux of 0.9 e^-^/Å^2^/s. Applied underfocus values ranged between 1 and 2.2 µm. The pixel size was calibrated to 0.935 Å as described before^34^. Details of data acquisition are summed up in Supplementary Table 1.

### Cryo-EM image processing and helical reconstruction

For all polymorphs, MotionCor2^60^ was used for movie correction and CTF parameters were fitted with CTFFIND4^61^. Fibrils were manually picked, and segments were extracted with an interbox distance of 10 % of the box sizes. Box sizes were chosen as 220 Å, 200 Å and 220 Å for PM1, PM2 and PM3 respectively. Further image processing, including 3D reconstructions, was done with RELION 3.0.5^62,63^. Gold-standard refinements were performed as described before^33^ by selecting entire fibrils and splitting the data set accordingly into an even and odd set. The Fourier shell correlation was computed between two half maps. According to the 0.143 criterion the obtained resolutions were 4.2 Å (PM1), 4.2 Å (PM2), and 8.1 Å (PM3) (Supplementary Figures 4, 7, and 8, respectively, for FSC curves). Image processing and reconstruction details for all polymorphs can be found in Supplementary Tables 2 and 3.

### Model building and refinement of polymorph 1

For PM1, a single chain atomic model was built with Coot^64,65^ by placing a poly-alanine model de novo into the density. After manual optimization of the protein backbone, sidechains were added and rotamers manually refined with respect to Ramachandran outliers and potential clashes. Five copies of the final single chain model were placed into the EM density map. The final model, containing six symmetry-related monomers of IAPP PM1, was used for real space refinement in PHENIX^66^ with manually assigned β-sheets. Subsequently, the model was refined by multiple rounds of optimization in Coot, PHENIX and MDFF^67,68^. MDFF was performed using an explicit solvent. The structure was embedded in a box of water and ions were added to the system (concentration 1.5 M). Secondary structure restraints, cispeptide as well as chirality restraints were applied. The scaling factor of the map potential was set to g = 0.3 and a time period of 10 ns was simulated. The final model of PM1 was obtained by averaging the coordinates of the MDFF trajectory and a final energy minimization with NCS restraints and position restraints using CNS^69,70^ including hydrogen atoms. B-factors were assigned based on RMSF values calculated from the MDFF trajectory. Model evaluation was done using MolProbity^71^. The final statistics of the refinement are shown in Supplementary Table 2. Molecular graphics and further analyses were performed using Chimera^72^ and ChimeraX^73^.

### Model building and refinement of polymorph 2

Because of difficulties assigning residues to the density of polymorph 2, two poly-alanine backbones each containing 21 residues were built both in forward and backward trace directions in Coot^64,65^. There is a total of 17 possible assignments of segments from the IAPP sequence to the 21 residues visible in the density. Accordingly, we performed 17 sidechain assignments for each backbone using Scwrl4^74^. The resulting 34 models were energy minimized with CNS^69^ and refined into the density map using DireX^35^. The C_free_ value^36^ is the real-space map correlation coefficient computed from the density map filtered with a band-pass from 3.0 to 4.0 Å. It served as a criterion to rank the models, as shown in Figure 5 and Supplementary Figure 6. The model that scored best according to this ranking was further refined using MDFF^67,68^. MDFF was performed with the same settings as for PM1. Refinement was finalized by averaging the coordinates of the MDFF trajectory.

### Molecular dynamics simulation

Molecular dynamics simulations were performed for the model of PM1 to test its stability. The starting structure for the simulation was built using CHARMM-GUI solution builder^75,76^ by inserting the cryo-EM structure of PM1 into a cubic water box containing 38907 water molecules and further added 10 chloride ions to neutralize the system. We have carried out two independent all-atom simulations using GROMACS^77^ (version 2019.3) and CHARMM36 force fields for protein^78^, water^79^ and ions^80^. The systems were first minimized using steepest descent algorithm in 5000 steps to remove the bad contacts followed by 500 ps (time step 1 fs) of equilibration in a constant volume and temperature ensemble. Later, two production runs of 250 ns were carried out under constant pressure and temperature conditions with a time-step of 2 fs by applying LINCS constraints to the bonds containing hydrogen atoms^81^. The temperature of systems maintained around 300 K using Nosé-Hoover thermostat^82,83^ and a pressure of 1 bar maintained using Parrinello-Rahman barostat^84^. Short-range electrostatic and van der Waals interactions were computed up to a cutoff of 12 Å using potential-shift and force-switch methods, respectively. Long-range electrostatic interactions beyond the 12 Å cutoff were computed using particle-mesh Ewald algorithm^85^.

## Acknowledgements

We thank P.J. Peters and C. López-Iglesias for advice and helpful discussions, H. Duimel for help with sample preparation, and the M4I Division of Nanoscopy of Maastricht University for microscope access and support. The authors gratefully acknowledge the computing time granted by the Jülich Aachen Research Alliance High-Peformance Computing (JARA-HPC) Vergabegremium and VSR commission on the supercomputer JURECA at Forschungszentrum Jülich. Support from a European Research Council (ERC) Consolidator Grant (grant agreement no. 726368) to W.H. is acknowledged. K.R.P. and G.F.S. acknowledge support from Helmholtz Association Initiative and Networking Fund under project number ZT-I-0003.

The authors declare no competing financial interests.

## Additional Information

### Accession codes

The structure of IAPP PM1 has been deposited in the Protein Data Bank under accession code 6Y1A. The cryo-EM density maps have been deposited in the Electron Microscopy Data Bank under accession codes EMD-10669 (PM1), EMD-10670 (PM2), EMD-10671 (PM3).

### Author contributions

L.G., W.H., T.K. and G.F.S. conceived the study. T.K. performed and analyzed fibril preparation and AFM experiments. R.G.B.R. performed cryo-EM experiments and initial data analysis. C.R., T.K. and G.F.S. performed image processing and initial reconstruction. C.R. and G.F.S. performed reconstruction, model building and refinement. L.U.S., K.R.P. and G.F.S. performed molecular dynamics simulations and structure fitting. C.R., T.K., G.F.S., W.H., L.G, K.R.P. and L.U.S wrote the manuscript. D.W. and all other authors discussed results and commented on the manuscript.

## References

1. Opie, E. L. On the relation of chronic interstitial pancreatitis to the islands of langerhans and to diabetes mellitus. J. Exp. Med. 5, 397–428 (1901).

2. Jurgens, C. A. et al. β-Cell loss and β-cell apoptosis in human type 2 diabetes are related to islet amyloid deposition. Am. J. Pathol. 178, 2632–2640 (2011).

3. Westermark, P., Andersson, A. & Westermark, G. T. Islet amyloid polypeptide, islet amyloid, and diabetes mellitus. Physiol. Rev. 91, 795–826 (2011).

4. Wimalawansa, S. J. Amylin, calcitonin gene-related peptide, calcitonin, and adrenomedullin: A peptide superfamily. Crit. Rev. Neurobiol. 11, 167–239 (1997).

5. Akter, R. et al. Islet Amyloid Polypeptide: Structure, Function, and Pathophysiology. Journal of Diabetes Research (2016). doi:10.1155/2016/2798269

6. Mukherjee, A., Morales-Scheihing, D., Butler, P. C. & Soto, C. Type 2 diabetes as a protein misfolding disease. Trends Mol. Med. 21, 439–449 (2015).

7. Cao, P., Abedini, A. & Raleigh, D. P. Aggregation of islet amyloid polypeptide: From physical chemistry to cell biology. Curr. Opin. Struct. Biol. 23, 82–89 (2013).

8. Halban, P. A. et al. β-Cell failure in type 2 diabetes: Postulated mechanisms and prospects for prevention and treatment. J. Clin. Endocrinol. Metab. 99, 1983–1992 (2014).

9. Zraika, S. et al. Toxic oligomers and islet beta cell death: Guilty by association or convicted by circumstantial evidence? Diabetologia 53, 1046–1056 (2010).

10. Zhang, S. et al. The pathogenic mechanism of diabetes varies with the degree of overexpression and oligomerization of human amylin in the pancreatic islet β cells. FASEB J. 28, 5083–5096 (2014).

11. Masters, S. L. et al. Activation of the NLRP3 inflammasome by islet amyloid polypeptide provides a mechanism for enhanced IL-1β 2 in type 2 diabetes. Nat. Immunol. 11, 897–904 (2010).

12. Rivera, J. F. et al. Human-IAPP disrupts the autophagy/lysosomal pathway in pancreatic B-cells: Protective role of p62-positive cytoplasmic inclusions. Cell Death Differ. 18, 415–426 (2011).

13. Gupta, D. & Leahy, J. L. Islet amyloid and type 2 diabetes: Overproduction or inadequate clearance and detoxification? J. Clin. Invest. 124, 3292–3294 (2014).

14. Casas, S. et al. Impairment of the ubiquitin-proteasome pathway is a downstream endoplasmic reticulum stress response induced by extracellular human islet amyloid polypeptide and contributes to pancreatic β-cell apoptosis. Diabetes 56, 2284–94 (2007).

15. Hull, R. L. et al. Amyloid formation in human IAPP transgenic mouse islets and pancreas, and human pancreas, is not associated with endoplasmic reticulum stress. Diabetologia 52, 1102–11 (2009).

16. Janson, J., Ashley, R. H., Harrison, D., McIntyre, S. & Butler, P. C. The mechanism of islet amyloid polypeptide toxicity is membrane disruption by intermediate-sized toxic amyloid particles. Diabetes 48, 491–8 (1999).

17. Paulsson, J. F. et al. High plasma levels of islet amyloid polypeptide in young with new-onset of type 1 diabetes mellitus. PLoS ONE 9, e93053 (2014).

18. Martinez-Valbuena, I. et al. Interaction of amyloidogenic proteins in pancreatic β cells from subjects with synucleinopathies. Acta Neuropathol. 135, 877–886 (2018).

19. Oskarsson, M. E. et al. In vivo seeding and cross-seeding of localized amyloidosis: A molecular link between type 2 diabetes and Alzheimer disease. Am. J. Pathol. 185, 834–846 (2015).

20. Moreno-Gonzalez, I. et al. Molecular interaction between type 2 diabetes and Alzheimer’s disease through cross-seeding of protein misfolding. Mol. Psychiatry 9, 1327–1334 (2017).

21. O’Nuallain, B., Williams, A. D., Westermark, P. & Wetzel, R. Seeding Specificity in Amyloid Growth Induced by Heterologous Fibrils. J. Biol. Chem. 279, 17490–9 (2004).

22. Kajava, A. V., Aebi, U. & Steven, A. C. The parallel superpleated beta-structure as a model for amyloid fibrils of human amylin. J. Mol. Biol. 348, 247–52 (2005).

23. Luca, S., Yau, W. M., Leapman, R. & Tycko, R. Peptide conformation and supramolecular organization in amylin fibrils: Constraints from solid-state NMR. Biochemistry 46, 13505–22 (2007).

24. Wiltzius, J. J. W. et al. Atomic structure of the cross-β spine of islet amyloid polypeptide (amylin). Protein Sci. 17, (2008).

25. Bedrood, S. et al. Fibril structure of human islet amyloid polypeptide. J. Biol. Chem. 287, 5235–41 (2012).

26. Alexandrescu, A. T. Amide Proton Solvent Protection in Amylin Fibrils Probed by Quenched Hydrogen Exchange NMR. PLoS ONE 8, :e56467 (2013).

27. Weirich, F. et al. Structural characterization of fibrils from recombinant human islet amyloid polypeptide by solid-state NMR: The central FGAILS segment is part of the β-sheet core. PLoS ONE 11, e0161243 (2016).

28. Goldsbury, C. S. et al. Polymorphic fibrillar assembly of human amylin. J. Struct. Biol. 119, 17–27 (1997).

29. Hutton, J. C. The internal pH and membrane potential of the insulin-secretory granule. Biochem. J. 204, 171–8 (1982).

30. Westermark, P., Engstrom, U., Johnson, K. H., Westermark, G. T. & Betsholtz, C. Islet amyloid polypeptide: Pinpointing amino acid residues linked to amyloid fibril formation. PNAS 87, 5036–40 (1990).

31. Betsholtz, C. et al. Sequence divergence in a specific region of islet amyloid polypeptide (IAPP) explains differences in islet amyloid formation between species. FEBS Lett. 251, 261–4 (1989).

32. Tenidis, K. et al. Identification of a penta- and hexapeptide of islet amyloid polypeptide (IAPP) with amyloidogenic and cytotoxic properties. J. Mol. Biol. 295, 1055–71 (2000).

33. Gremer, L. et al. Fibril structure of amyloid-β(1–42) by cryo–electron microscopy. Science 358, 116–119 (2017).

34. Röder, C. et al. Atomic structure of PI3-kinase SH3 amyloid fibrils by cryo-electron microscopy. Nat. Commun. 10, 3754 (2019).

35. Wang, Z. & Schröder, G. F. Real-space refinement with DireX: From global fitting to side-chain improvements. Biopolymers 97, 687–97 (2012).

36. Falkner, B. & Schröder, G. F. Cross-validation in cryo-EM-based structural modeling. PNAS 110, 8930–8935 (2013).

37. Padrick, S. B. & Miranker, A. D. Islet amyloid polypeptide: Identification of long-range contacts and local order on the fibrillogenesis pathway. J. Mol. Biol. 308, 783–94 (2001).

38. Chen, M. S. et al. Characterizing the assembly behaviors of human amylin: A perspective derived from C-terminal variants. Chem. Commun. 49, 1799–801 (2013).

39. Yonemoto, I. T., Kroon, G. J. A., Dyson, H. J., Balch, W. E. & Kelly, J. W. Amylin proprotein processing generates progressively more amyloidogenic peptides that initially sample the helical state. Biochemistry 47, 9900–10 (2008).

40. Sakagashira, S. et al. Missense mutation of amylin gene (S20G) in Japanese NIDDM patients. Diabetes 45, 1279–81 (1996).

41. Seino, S. S20G mutation of the amylin gene is associated with Type II diabetes in Japanese. Diabetologia 44, 906–9 (2001).

42. Meier, D. T. et al. The S20G substitution in hIAPP is more amyloidogenic and cytotoxic than wild-type hIAPP in mouse islets. Diabetologia 59, 2166–71 (2016).

43. Cao, P. et al. Sensitivity of amyloid formation by human islet amyloid polypeptide to mutations at residue 20. J. Mol. Biol. 421, 282–95 (2012).

44. Sakagashira, S. et al. S20G mutant amylin exhibits increased in vitro amyloidogenicity and increased intracellular cytotoxicity compared to wild-type amylin. Am. J. Pathol. 157, 2101–9 (2000).

45. Ma, Z. et al. Enhanced in vitro production of amyloid-like fibrils from mutant (s20g) islet amyloid polypeptide. Amyloid 8, 242–9 (2001).

46. Xu, W., Jiang, P. & Mu, Y. Conformation preorganization: Effects of S20G mutation on the structure of human islet amyloid polypeptide segment. J. Phys. Chem. B 113, 7308–14 (2009).

47. Mirecka, E. A. et al. β-Hairpin of Islet Amyloid Polypeptide Bound to an Aggregation Inhibitor. Sci. Rep. 6, (2016).

48. Nilsberth, C. et al. The ‘Arctic’ APP mutation (E693G) causes Alzheimer’s disease by enhanced Aβ protofibril formation. Nat. Neurosci. (2001). doi:10.1038/nn0901-887

49. Kruger, D. F. & Gloster, M. A. Pramlintide for the treatment of insulin-requiring diabetes mellitus: Rationale and review of clinical data. Drugs 64, 1419–32 (2004).

50. Roth, J. D. et al. Leptin responsiveness restored by amylin agonism in diet-induced obesity: Evidence from nonclinical and clinical studies. PNAS 105, 7257–62 (2008).

51. Wang, H., Abedini, A., Ruzsicska, B. & Raleigh, D. P. Rationally Designed, Nontoxic, Nonamyloidogenic Analogues of Human Islet Amyloid Polypeptide with Improved Solubility. Biochemistry 53, 5876–84 (2014).

52. Janson, J. et al. Increased Risk of Type 2 Diabetes in Alzheimer Disease. Diabetes 53, 474–81 (2004).

53. Yang, Y. & Song, W. Molecular links between Alzheimer’s disease and diabetes mellitus. Neuroscience 250, 140–50 (2013).

54. Colvin, M. T. et al. Atomic Resolution Structure of Monomorphic Aβ42 Amyloid Fibrils. J. Am. Chem. Soc. (2016). doi:10.1021/jacs.6b05129

55. Wälti, M. A. et al. Atomic-resolution structure of a disease-relevant Aβ(1-42) amyloid fibril. PNAS 113, 4976–84 (2016).

56. Tycko, R. Molecular Structure of Aggregated Amyloid-β: Insights from Solid State Nuclear Magnetic Resonance. Q. Rev. Biophys. 39, 1–55 (2016).

57. Xiao, Y. et al. Aβ(1-42) fibril structure illuminates self-recognition and replication of amyloid in Alzheimer’s disease. Nat. Struct. Mol. Biol. (2015). doi:10.1038/nsmb.2991

58. Kollmer, M. et al. Cryo-EM structure and polymorphism of Aβ amyloid fibrils purified from Alzheimer’s brain tissue. Nat. Commun. (2019). doi:10.1038/s41467-019-12683-8

59. Colvin, M. T. et al. Atomic resolution structure of monomorphic Aβ(42) amyloid fibrils. J. Am. Chem. Soc. 138, 9663–9674 (2016).

60. Zheng, S. Q., Palovcak, E., Armache, J.-P., Cheng, Y. & Agard, D. A. MotionCor2: Anisotropic correction of beam-induced motion for improved single-particle electron cryo-microscopy. Nat. Methods 14, 331–332 (2017).

61. Rohou, A. & Grigorieff, N. CTFFIND4: Fast and accurate defocus estimation from electron micrographs. J Struct Biol 192, 216–221 (2015).

62. He, S. & Scheres, S. H. W. Helical reconstruction in RELION. J. Struct. Biol. 198, 163–176 (2017).

63. Scheres, S. H. W. RELION: Implementation of a Bayesian approach to cryo-EM structure determination. J. Struct. Biol. 180, 519–530 (2012).

64. Emsley, P. & Cowtan, K. Coot: Model-building tools for molecular graphics. Acta Crystallogr. Sect. D Biol. Crystallogr. 60, 2126–2132 (2004).

65. Emsley, P., Lohkamp, B., Scott, W. G. & Cowtan, K. Features and development of Coot. Acta Crystallogr. Sect. D Biol. Crystallogr. 66, 486–501 (2010).

66. Adams, P. D. et al. PHENIX: A comprehensive Python-based system for macromolecular structure solution. Acta Crystallogr. Sect. D Biol. Crystallogr. 66, 213–221 (2010).

67. Trabuco, L. G., Villa, E., Schreiner, E., Harrison, C. B. & Schulten, K. Molecular dynamics flexible fitting: A practical guide to combine cryo-electron microscopy and X-ray crystallography. Methods 49, 174–180 (2009).

68. Trabuco, L. G., Villa, E., Mitra, K., Frank, J. & Schulten, K. Flexible Fitting of Atomic Structures into Electron Microscopy Maps Using Molecular Dynamics. Structure 16, 673–683 (2008).

69. Brunger, A. T. et al. Crystallography & NMR system: A new software suite for macromolecular structure determination. Acta Crystallogr. Sect. D Biol. Crystallogr. 54, 905–921 (1998).

70. Brunger, A. T. Version 1.2 of the crystallography and nmr system. Nat. Protoc. 2, 2728–2733 (2007).

71. Chen, V. B. et al. MolProbity: All-atom structure validation for macromolecular crystallography. Acta Crystallogr. Sect. D Biol. Crystallogr. (2010). doi:10.1107/S0907444909042073

72. Pettersen, E. F. et al. UCSF Chimera - A visualization system for exploratory research and analysis. J. Comput. Chem. 25, 1605–1612 (2004).

73. Goddard, T. D. et al. UCSF ChimeraX: Meeting modern challenges in visualization and analysis. Protein Sci. 27, 14–25 (2018).

74. Krivov, G. G., Shapovalov, M. V. & Dunbrack, R. L. Improved prediction of protein side-chain conformations with SCWRL4. Proteins Struct. Funct. Bioinforma. (2009). doi:10.1002/prot.22488

75. Jo, S., Kim, T., Iyer, V. G. & Im, W. CHARMM-GUI: A web-based graphical user interface for CHARMM. J. Comput. Chem. (2008). doi:10.1002/jcc.20945

76. Lee, J. et al. CHARMM-GUI Input Generator for NAMD, GROMACS, AMBER, OpenMM, and CHARMM/OpenMM Simulations Using the CHARMM36 Additive Force Field. J. Chem. Theory Comput. 12, 405–413 (2016).

77. Abraham, M. J. et al. GROMACS: High performance molecular simulations through multi-level parallelism from laptops to supercomputers. SoftwareX 1–2, 19–25 (2015).

78. Best, R. B. et al. Optimization of the additive CHARMM all-atom protein force field targeting improved sampling of the backbone φ, ψ and side-chain χ^1^ and χ^2^ Dihedral Angles. J. Chem. Theory Comput. 8, 3256–3237 (2012).

79. MacKerell, A. D. et al. All-atom empirical potential for molecular modeling and dynamics studies of proteins. J. Phys. Chem. B 102, 3586–3616 (1998).

80. Beglov, D. & Roux, B. Finite representation of an infinite bulk system: Solvent boundary potential for computer simulations. J. Chem. Phys. 100, 9050–9063 (1994).

81. Hess, B., Bekker, H., Berendsen, H. J. C. & Fraaije, J. G. E. M. LINCS: A Linear Constraint Solver for molecular simulations. J. Comput. Chem. 18, 1463–72 (1997).

82. Nosé, S. A unified formulation of the constant temperature molecular dynamics methods. J. Chem. Phys. 81, 511–519 (1984).

83. Hoover, W. G. Canonical dynamics: Equilibrium phase-space distributions. Phys. Rev. A 31, 1695–1697 (1985).

84. Parrinello, M. & Rahman, A. Polymorphic transitions in single crystals: A new molecular dynamics method. J. Appl. Phys. 52, 7182–7190 (1981).

85. Darden, T., York, D. & Pedersen, L. Particle mesh Ewald: An N·log(N) method for Ewald sums in large systems. J. Chem. Phys. 98, 10089–10092 (1993).

