## Supplementary Material for "Amyloid fibril structure of islet amyloid polypeptide by cryo-electron microscopy reveals similarities with amyloid beta"

### Supplementary Tables

#### Supplementary Table 1: Data collection for IAPP

| **Data Collection** | |
|---|---|
| Microscope | Tecnai Arctica |
| Camera | Falcon 3 (counting mode) |
| Acceleration voltage (kV) | 200 |
| Magnification | 110,000 |
| Defocus range (µm) | -1.0 to -2.2 |
| Dose rate ($e^{-}$/Å$^{2}$/s) | 0.9 |
| Exposure time (s) | 46 |
| Number of movie frames | 1800 |
| Pixel size (Å) | 0.935 |

#### Supplementary Table 2: Reconstruction, modeling and refinement of polymorph 1 (PM1).

| | **Polymorph 1** |
|---|---|
| **Reconstruction** | |
| Box size (pixel) | 220 |
| No. fibrils in final reconstruction | 1161 |
| No. segments in final reconstruction | 37,120 |
| Resolution (0.143 criterion) (Å) | 4.2 |
| Helical rise (Å) | 2.351 |
| Helical twist (°) | 178.23 |
| Helical pitch (Å) | 479.5 |
| Symmetry | C1 |
| **Model Composition** | |
| Non-hydrogen atoms | 2975 |
| Number of chains | 16 |
| **Model Refinement** | |
| Map CC (around atoms) | 0.9018 |
| RMSD bonds (Å) | 0.0039 |
| RMSD angles (°) | 0.60 |
| All-atom clash score | 15.2 |
| Ramachandran outliers/favored (%) | 0 / 95.7 |
| Rotamer outliers (%) | 0 |
| C-beta deviations | 0 |
| Molprobity score | 1.99 |

**Supplementary Table 3: Reconstruction of polymorphs 2 (PM2) and 3 (PM3).**

| | **Polymorph 2** | **Polymorph 3** |
|---|---|---|
| **Reconstruction** | | |
| Box size (pixel) | 200 | 220 |
| No fibrils in final reconstruction | 1480 | 99 |
| No segments in final reconstruction | 24011 | 4591 |
| Resolution (0.143 criterion) (Å) | 4.2 | 8.1 |
| Helical rise (Å) | 2.352 | 2.323 |
| Helical twist (°) | 179.10 | 179.47 |
| Helical pitch (Å) | 940 | 1590 |
| Symmetry | C1 (approx. $2_1$ screw) | C1 (approx. $2_1$ screw) |

### Supplementary Figures

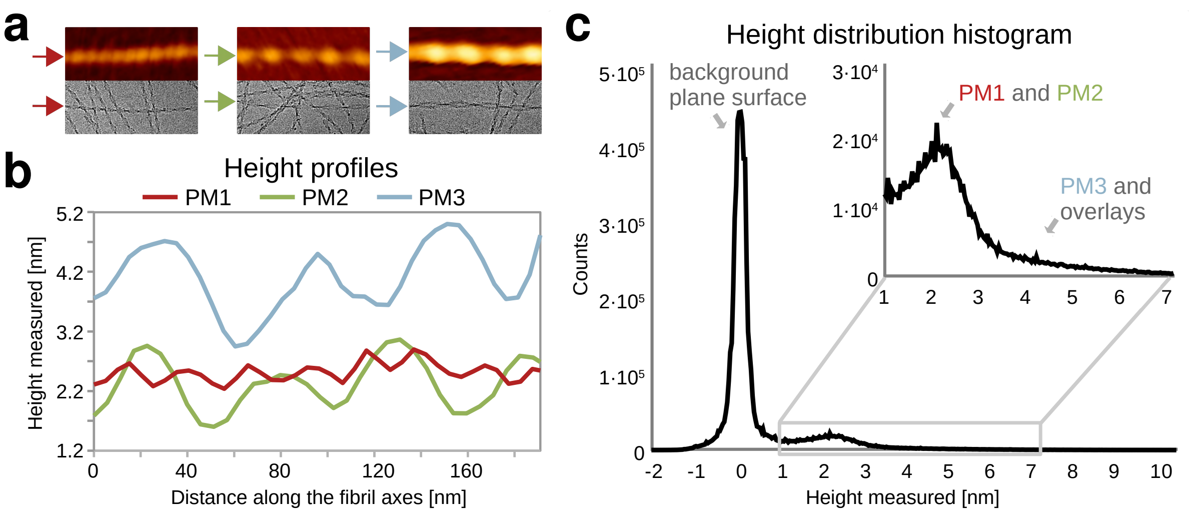


**Supplementary Figure 1: Comparison of described IAPP polymorphs. a** Single fibril cut-outs of polymorphs PM1, PM2 and PM3 from AFM images (top row) and cryo-EM micrographs (bottom row); single box size is 100x250 nm. **b** Height profiles of individual fibrils extracted from AFM images. **c** Height distribution histogram, showing the highest number of counts for the plane background surface around 0 nm and a distinct peak around 2.2 nm. The peak around 2.2 nm includes both PM1 and PM2 which are non-distinguishable in sense of height distribution. Moreover, a pronounced shoulder on the right corresponds to the presence of lower amounts of PM3 as well as the overlaps of single PM1/PM2 fibrils. For the height distribution analysis, histograms from six height images of 5x5 μm size and a resolution of 1024x1024 pixels were obtained, binned and presented in one graph. An example of the image used can be seen in Supplementary Figure 2.

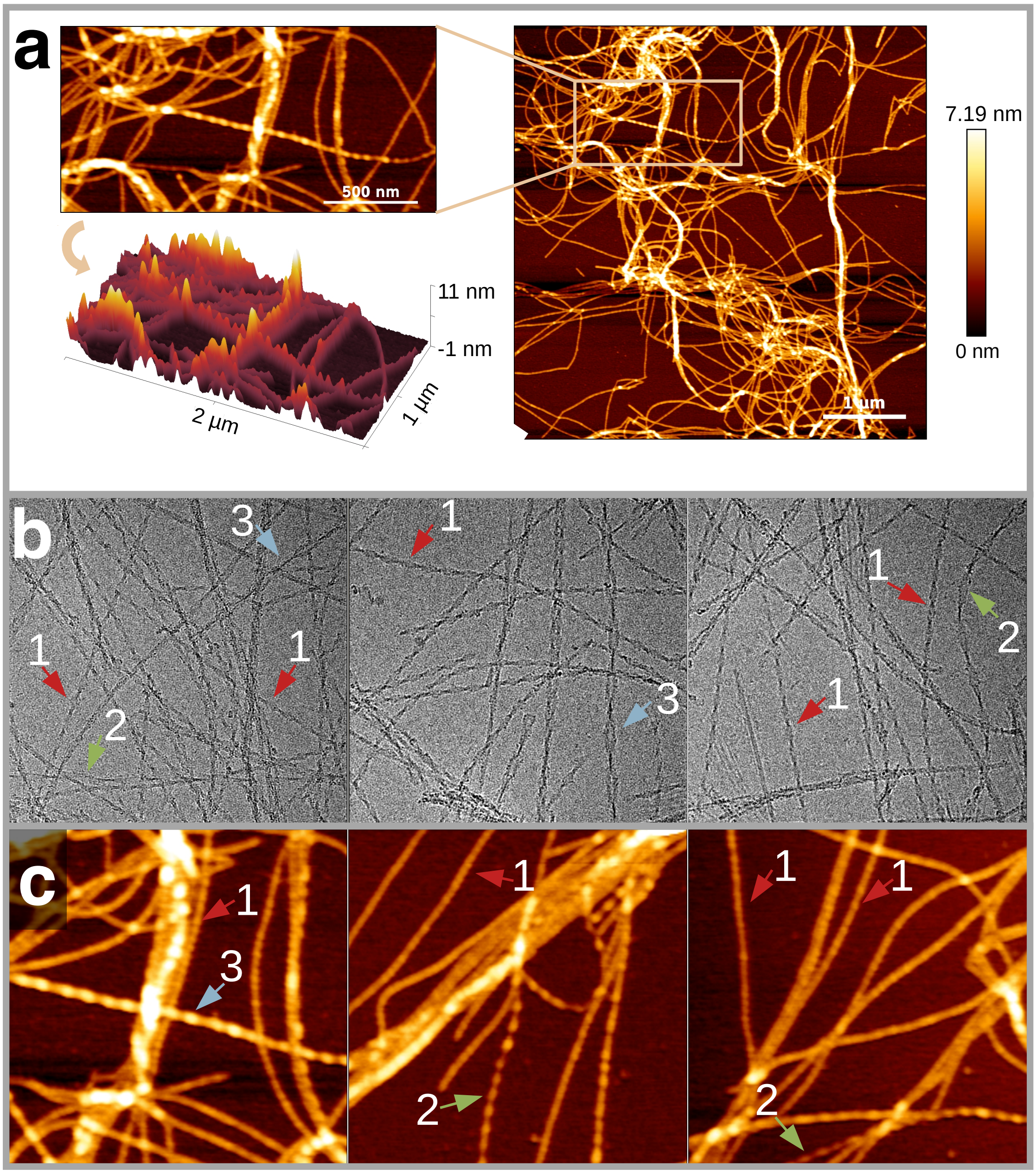


**Supplementary Figure 2: Overview of IAPP polymorphs**. **a** Typical height profile AFM image used for polymorph distribution analysis. **b** Cryo-EM micrographs showing 370x370 nm areas. **c** AFM overview images showing 1x1 µm areas. Arrows indicate the presence of PM1 (red), PM2 (green) and PM3 (blue).

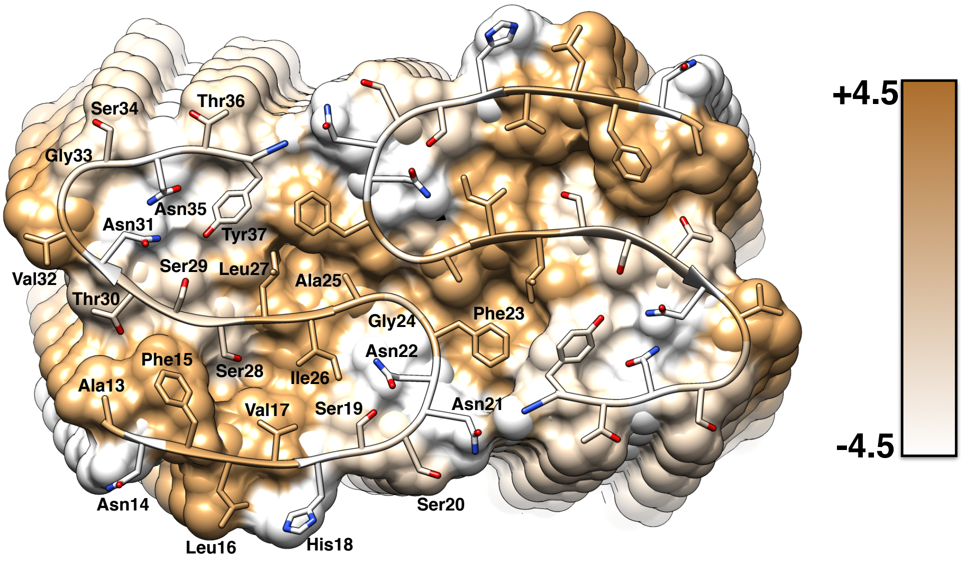


**Supplementary Figure 3: Hydrophobicity plot of the fibril displayed as top view**. Hydrophobicity levels of the IAPP polymorph 1 (PM1) fibril are colored according to Kyte-Doolittle[1] in the hydrophobicity score range -4.5 (white) to 4.5 (gold). One hydrophobic cluster spans the entire diagonal of the fibril cross-section. This hydrophobic streak is surrounded by highly ordered polar clusters.

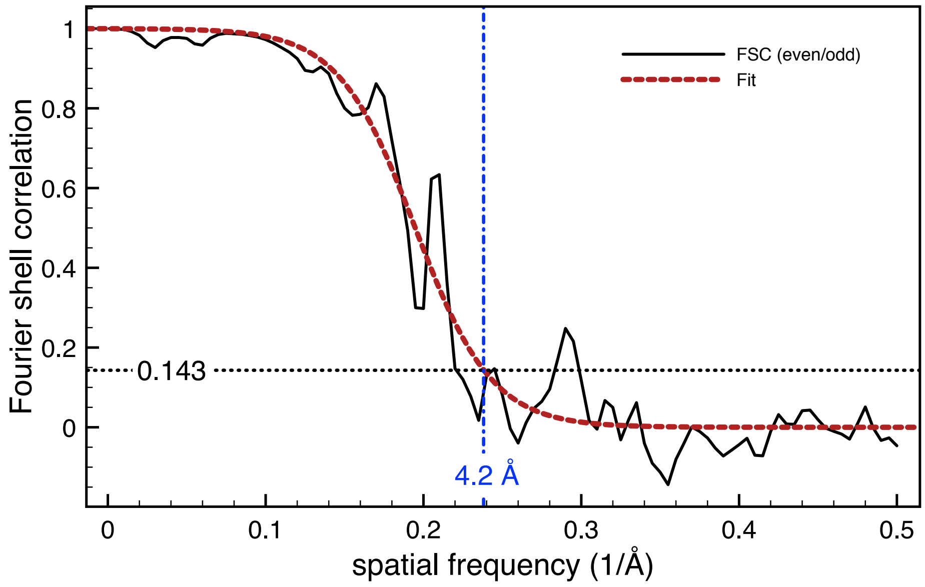


**Supplementary Figure 4: FSC Analysis of polymorph 1 (PM1).** FSC curves from the even/odd test (solid black) from the gold-standard refinement yields a resolution of 4.2 Å (using the 0.143 criterion). The even/odd FSC curve is fitted (red) with the model function 1/( 1+exp((x-A)/B) ) (with A=0.1947 and B=0.026) to obtain a more robust resolution estimate.

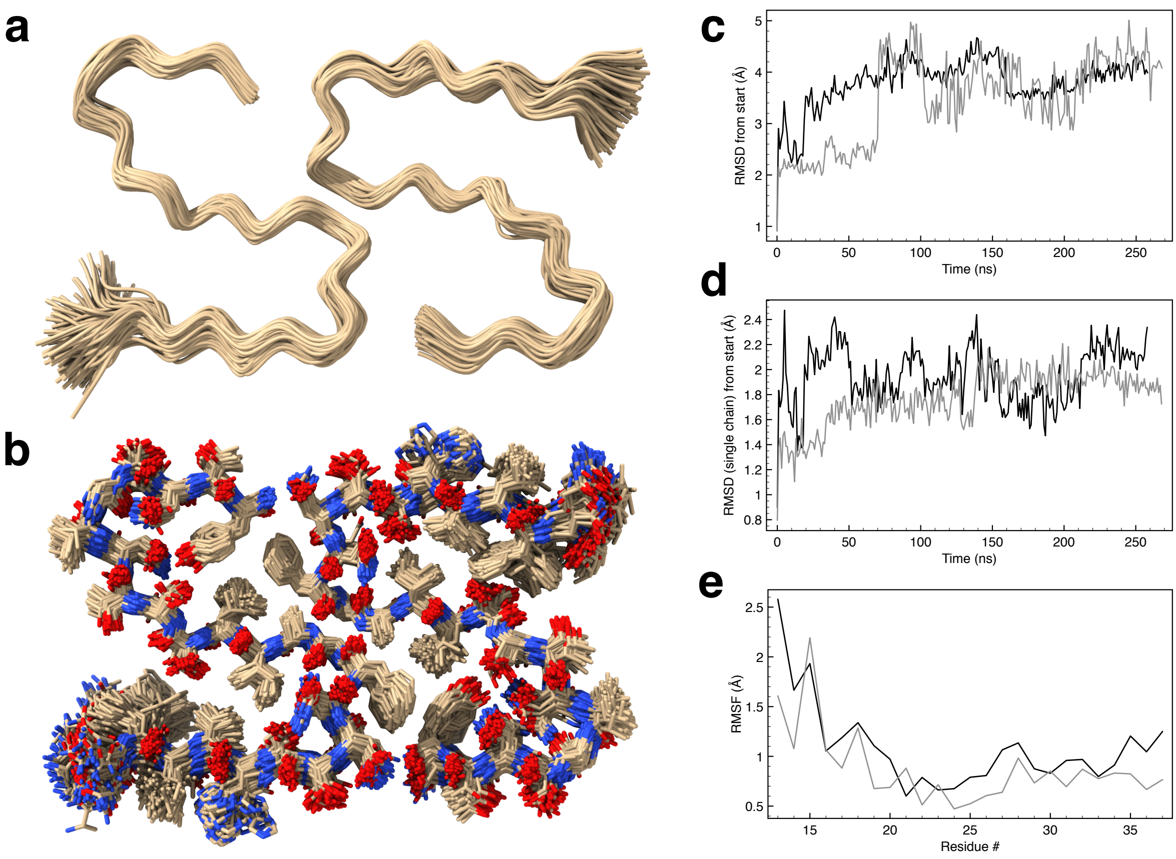


**Supplementary Figure 5: Results of molecular dynamics simulations of IAPP polymorph 1 (PM1).** Superimposed snapshots from a 250 ns simulation displaying only the backbone (**a**) or all atoms (except for solvent and hydrogen) (**b**). **c** Showing the RMSD from the deposited structure of PM1 (PDB ID 6Y1A) for two 250 ns simulations (black and grey lines, respectively). **d** Showing the RMSD of a single chain from the deposited structure during the two 250 ns simulations. **e** Showing the atomic root mean square fluctuations (RMSF) for each residue calculated over each 250 ns simulation.

| | Cwork | Cfree | | 1 5 10 15 20 | |
|---|---|---|---|---|---|
| 1 | 0.845 | 0.180 | KCNTATCATQRLAN | FLVHSSNNFGAILSSTNVGSN | TY |
| 2 | 0.807 | 0.151 | KC | NTATCATQRLANFLVHSSNNF | GAILSSTNVGSNTY |
| 3 | 0.818 | 0.133 | KCNTATCATQRLA | NFLVHSSNNFGAILSSTNVGS | NTY |
| 4 | 0.828 | 0.128 | KCNTATCATQR | LANFLVHSSNNFGAILSSTNV | GSNTY |
| 5 | 0.815 | 0.125 | KCNTATCATQRLANF | LVHSSNNFGAILSSTNVGSNT | Y |
| 6 | 0.798 | 0.124 | KCNTATC | ATQRLANFLVHSSNNFGAILS | STNVGSNTY |
| 7 | 0.768 | 0.121 | YTNS | GVNTSSLIAGFNNSSHVLFNA | LRQTACTATNCK |
| 8 | 0.755 | 0.118 | YTNSGVN | TSSLIAGFNNSSHVLFNALRQ | TACTATNCK |
| 9 | 0.789 | 0.115 | YTNSGVNT | SSLIAGFNNSSHVLFNALRQT | ACTATNCK |
| 10 | 0.832 | 0.111 | KCNTATCATQ | RLANFLVHSSNNFGAILSSTN | VGSNTY |
| 11 | 0.798 | 0.110 | KCNTATCATQRLANFL | VHSSNNFGAILSSTNVGSNTY | |
| 12 | 0.811 | 0.109 | YTNSGVNTSSL | IAGFNNSSHVLFNALRQTACT | ATNCK |
| 13 | 0.796 | 0.101 | KCNTATCA | TQRLANFLVHSSNNFGAILSS | TNVGSNTY |
| 14 | 0.751 | 0.095 | Y | TNSGVNTSSLIAGFNNSSHVL | FNALRQTACTATNCK |
| 15 | 0.771 | 0.095 | YTNSGV | NTSSLIAGFNNSSHVLFNALR | QTACTATNCK |
| 16 | 0.780 | 0.094 | YTN | SGVNTSSLIAGFNNSSHVLFN | ALRQTACTATNCK |
| 17 | 0.813 | 0.094 | KCN | TATCATQRLANFLVHSSNNFG | AILSSTNVGSNTY |
| 18 | 0.817 | 0.090 | KCNTA | TCATQRLANFLVHSSNNFGAI | LSSTNVGSNTY |
| 19 | 0.763 | 0.085 | YT | NSGVNTSSLIAGFNNSSHVLF | NALRQTACTATNCK |
| 20 | 0.807 | 0.082 | KCNTATCAT | QRLANFLVHSSNNFGAILSST | NVGSNTY |
| 21 | 0.799 | 0.079 | KCNT | ATCATQRLANFLVHSSNNFGA | ILSSTNVGSNTY |
| 22 | 0.781 | 0.072 | YTNSGVNTSSLIA | GFNNSSHVLFNALRQTACTAT | NCK |
| 23 | 0.779 | 0.071 | YTNSGVNTSSLIAGFN | NSSHVLFNALRQTACTATNCK | |
| 24 | 0.766 | 0.071 | | YTNSGVNTSSLIAGFNNSSHV | LFNALRQTACTATNCK |
| 25 | 0.795 | 0.069 | K | CNTATCATQRLANFLVHSSNN | FGAILSSTNVGSNTY |
| 26 | 0.758 | 0.063 | YTNSGVNTS | SLIAGFNNSSHVLFNALRQTA | CTATNCK |
| 27 | 0.825 | 0.058 | KCNTAT | CATQRLANFLVHSSNNFGAIL | SSTNVGSNTY |
| 28 | 0.792 | 0.054 | KCNTATCATQRL | ANFLVHSSNNFGAILSSTNVG | SNTY |
| 29 | 0.800 | 0.054 | | KCNTATCATQRLANFLVHSSN | NFGAILSSTNVGSNTY |
| 30 | 0.763 | 0.050 | YTNSG | VNTSSLIAGFNNSSHVLFNAL | RQTACTATNCK |
| 31 | 0.743 | 0.045 | YTNSGVNTSSLIAG | FNNSSHVLFNALRQTACTATN | CK |
| 32 | 0.754 | 0.042 | YTNSGVNTSS | LIAGFNNSSHVLFNALRQTAC | TATNCK |
| 33 | 0.758 | 0.033 | YTNSGVNTSSLIAGF | NNSSHVLFNALRQTACTATNC | K |
| 34 | 0.757 | 0.025 | YTNSGVNTSSLI | AGFNNSSHVLFNALRQTACTA | TNCK |
| | | | | 1 5 10 15 20 | |

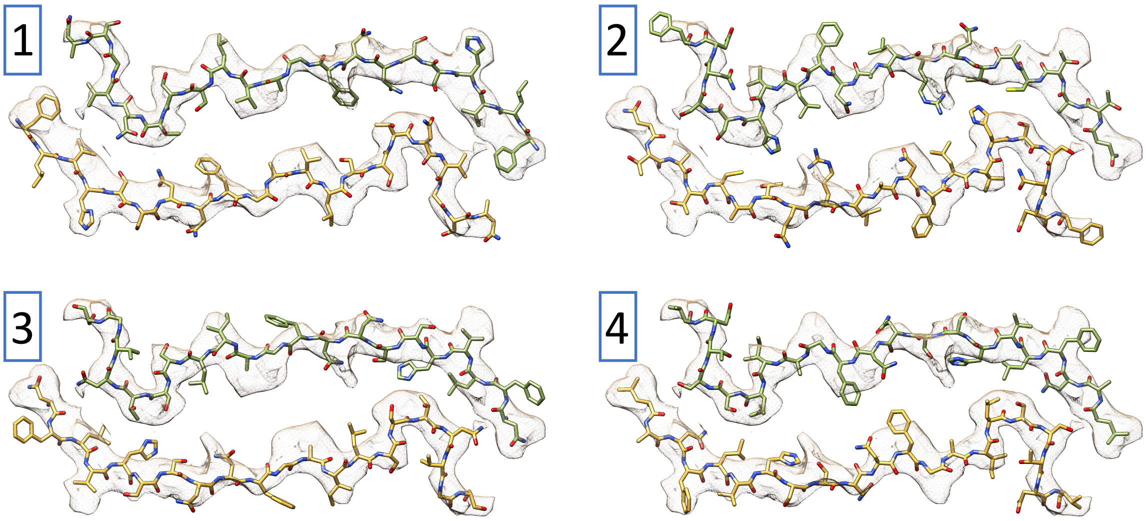


**Supplementary Figure 6: DireX² analysis of polymorph 2 (PM2)**. Ranking of the potential IAPP sequence snippets that have been modeled into the reconstructed density according to their $C_{free}$ values. Each snippet contains 21 amino acid residues which were modeled into the density map in both possible Cα-chain directions. Atomic models of the four most favorable sequence snippets are shown at the bottom. Note that some models, for example model 2, can be excluded since they are incompatible with the disulfide bond between residues $Cys^2$ and $Cys^7$.

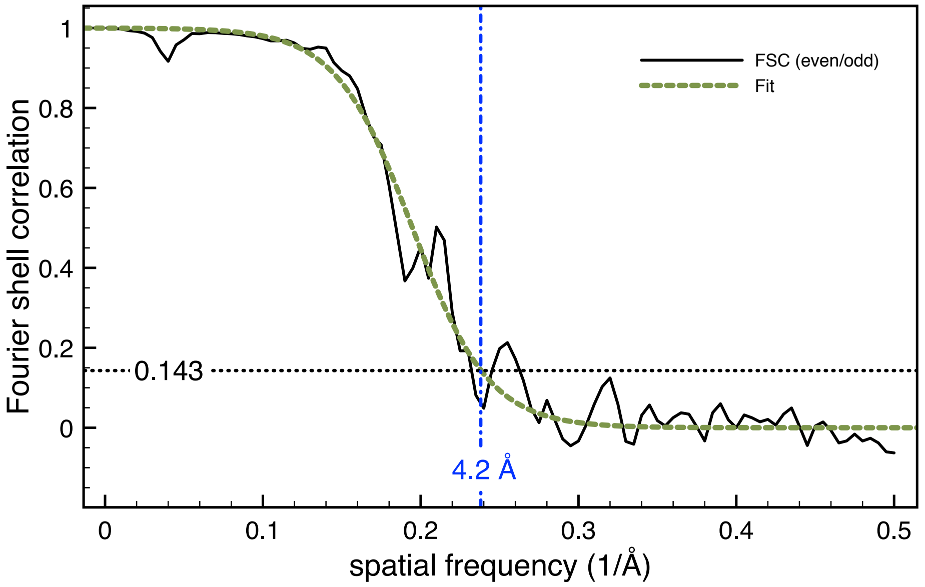


**Supplementary Figure: 7 FSC analysis of polymorph 2 (PM2).** FSC curves from the even/odd test (solid black) from the gold-standard refinement yields a resolution of 4.2 Å (using the 0.143 criterion). The even/odd FSC curve is fitted (green) with the model function 1/( 1+exp((x-A)/B)) ) (with A=0.194789 and B=0.02427) to obtain a more robust resolution estimate.

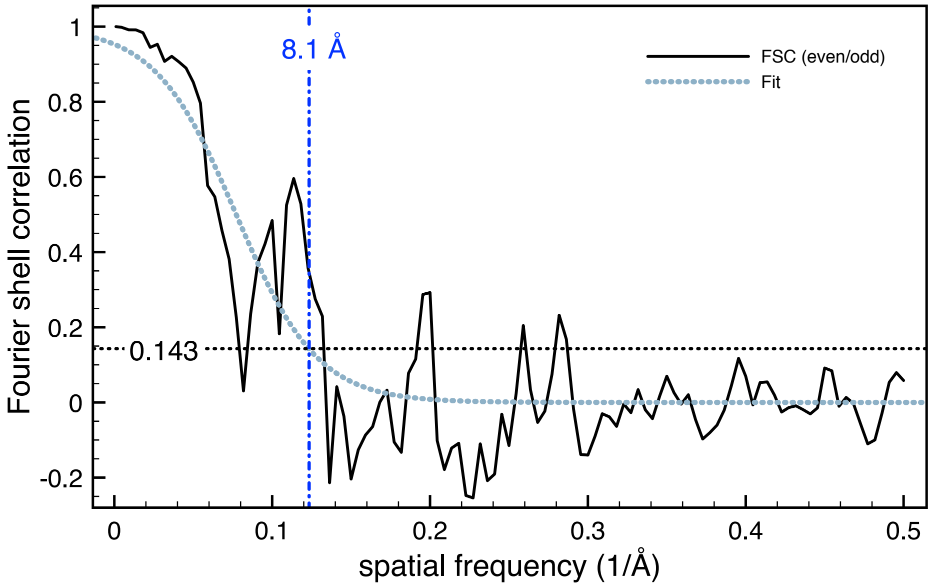


**Supplementary Figure 8: FSC analysis of Polymorph 3 (PM3).** FSC curves from the even/odd test (solid black) from the gold-standard refinement yields a resolution of 8.1 Å (using the 0.143 criterion). The even/odd FSC curve is fitted (light blue) with the model function 1/( 1+exp((x-A)/B) ) (with A=0.0772 and B=0.0256) to obtain a more robust resolution estimate.
